# Distinct Neuronal Populations Contribute to Trace Conditioning and Extinction Learning in the Hippocampal CA1

**DOI:** 10.1101/2020.03.06.980854

**Authors:** Rebecca A. Mount, Kyle R. Hansen, Sudiksha Sridhar, Ali I. Mohammed, Moona Abdulkerim, Robb Kessel, Bobak Nazer, Howard J. Gritton, Xue Han

## Abstract

Trace conditioning and extinction learning depend on the hippocampus, but it remains unclear how ongoing neural activities in the hippocampus are modulated during different learning processes. To explore this question, we performed calcium imaging in a large number of individual CA1 neurons during both trace eye-blink conditioning and subsequent extinction learning in mice. Using trial-averaged calcium fluorescence analysis, we found direct evidence that in real time, as learning emerges, distinct populations of CA1 cells contribute to trace conditioned learning versus extinction learning. Furthermore, we examined network connectivity by calculating co-activity between CA1 neuron pairs, and found that CA1 network connectivity is different between conditioning and extinction and between correct versus incorrect behavioral responses during trace conditioned learning. However, the overall connectivity density remains constant across these behavioral conditions. Together, our results demonstrate that distinct populations of CA1 neurons, forming different sub-networks with unique connectivity patterns, encode different aspects of learning.

## Introduction

The hippocampus is critical for learning and memory in animals and humans. Early surgical lesions of the hippocampus in human patients, designed to alleviate intractable epilepsy, resulted in severe memory loss and an inability to form new declarative or episodic memories^1,2^. Hippocampal atrophy is also associated with diseases related to memory loss and cognitive decline including dementia and Alzheimer’s disease^3–7^. Many mechanistic studies have highlighted the importance of the hippocampus for spatial, contextual, and associative learning in a variety of animal models^8,9^.

Various experimental paradigms have been devised to probe hippocampal-dependent forms of learning and memory. One such well-established paradigm is trace eye blink conditioning, which requires an intact hippocampus^10–12^. In this experimental design, subjects are presented with a conditioned stimulus (CS), such as a tone or light, which reliably predicts an unconditioned stimulus (US), such as a puff of air or electrical shock delivered to the subject’s eyelid. In trace conditioning, the CS and US are separated temporally by a quiescent trace interval. Over time, subjects will learn to associate the CS with the US, generating a behavioral conditioned response to the CS^13–18^. Trace conditioning acquisition depends on signaling at both nicotinic and muscarinic acetylcholine receptors (AChRs)^19–26^ and is mediated through NMDA receptor-dependent plasticity^27^.

The hippocampus is also required for context-dependent extinction learning^11^. Extinction learning is traditionally considered new learning that overrides a previously learned relationship. In the example of trace conditioning, the subject learns that the previously established CS is no longer predictive of a subsequent US. Extinction learning after trace conditioning can be tested by the presentation of the CS without the associated US, and monitoring the strength or presence of a conditioned response. As new learning occurs, subjects will suppress their conditioned response to the previously predictive tone or light. Extinction learning has also been shown to be NMDA receptor dependent^28^, and requires the involvement of hippocampal inhibitory neurons^29^ and adult neurogenesis^30^.

While the hippocampus is known to be important in both trace conditioning and extinction learning, it remains largely unknown how individual hippocampal neurons selectively participate in this learning. Two distinct functional populations related to fear conditioning and extinction have been reported in the amygdala^31^, and segregated populations of hippocampal CA1 neurons upregulate different genes for both fear conditioning and context-dependent fear extinction^32^, supporting the idea that trace conditioning and extinction learning involve distinct learning mechanisms encoded by different neurons. However, changes in CA1 populations were quantified at later time points, after learning occurred, leaving in question whether neural activity changes during learning, or is a result of plasticity changes in the minutes to hours after new learning. In this study, we sought to measure the ongoing neuronal activity of individual neurons during trace eye-blink conditioning and subsequent extinction learning to better understand the time course and mechanics of how these two types of learning might interact in the neural population.

In order to address these questions, we performed calcium imaging of individual CA1 neurons in mice over multiple days during the course of trace eye-blink conditioning. Calcium imaging allows us to measure hundreds to thousands of neurons simultaneously with single-cell resolution, across multiple trials and multiple days of learning^33,34^. Once conditioning was achieved, mice underwent a final conditioning session followed by extinction training, enabling us to track the same neuron population during both learning paradigms. Hippocampal-dependent trace conditioning is a learning task well suited to calcium imaging because learning, and the associated CA1 neuronal responses, evolves gradually, unlike fear conditioning, where learning can occur as rapidly as a single trial.

Using trial-averaged approaches we found that a significant fraction of CA1 neurons showed CS-related responses for both trace conditioning and extinction learning. However, the identities of the cells differed between trace conditioning and extinction learning, suggesting two functionally distinct sub-populations of cells within the hippocampus CA1. To further understand how the neural populations reflect learning as it is occurring, we analyzed co-activity between CA1 neuron pairs on a trial-by-trial basis. Using this trial-by-trial analysis method, we found that populations of neuron pairs are differentially activated during trace conditioning versus extinction learning. In addition, this method revealed that during trace conditioning, neurons are also differentially co-active on trials during which the animal exhibited the “correct” versus the “incorrect” behavioral response, highlighting the potential of trial-by-trial co-activity analysis to detect features of network response.

## Results

### Conditioned responding increases across trace-conditioning sessions in a classical eye blink task and decreases during extinction

Trace conditioning experiments were performed in head-fixed mice (n=9 mice) that were positioned under a custom-built wide-field microscope (Figure 1A) equipped with a scientific (sCMOS) camera, as previously described^33^. Calcium activity in CA1 neurons was monitored via GCaMP6f fluorescence, which allows recording from hundreds of neurons simultaneously^35–40^ (Figure 1A). Prior to imaging, mice were injected with AAV-Syn-GCaMP6f and implanted with a custom window that allowed optical access to dorsal CA1 (Figure 1C). 4-6 weeks after surgery, mice were habituated and then trained on a classic trace eye-blink conditioning paradigm followed by an extinction training session (Figure 1B). The paradigm consisted of a 9500Hz, 350ms tone as a conditioned stimulus (CS), followed by a 250ms trace interval, followed by a 100ms gentle puff of air to one eye that served as the unconditioned stimulus (US) (Figure 1D). Eye behavior was monitored with a USB 3.0 Camera (Figure 1A,Ei). Animals were trained for 60-80 CS-US trials over 5-9 days, until they reached conditioned response criterion (conditioned response on 65% of trials). After reliable conditioned response to CS presentations was established, on the final day of imaging animals were given a block of 20-40 CS-US trials (last training session), followed by a block of CS-only extinction trials, where the CS was not followed by the US (extinction session, Figure 1B). In this paradigm, we could perform calcium imaging of the same neurons during both learning conditions, allowing us to track how activity of each neuron changes during extinction acquisition.

**Figure 1.**
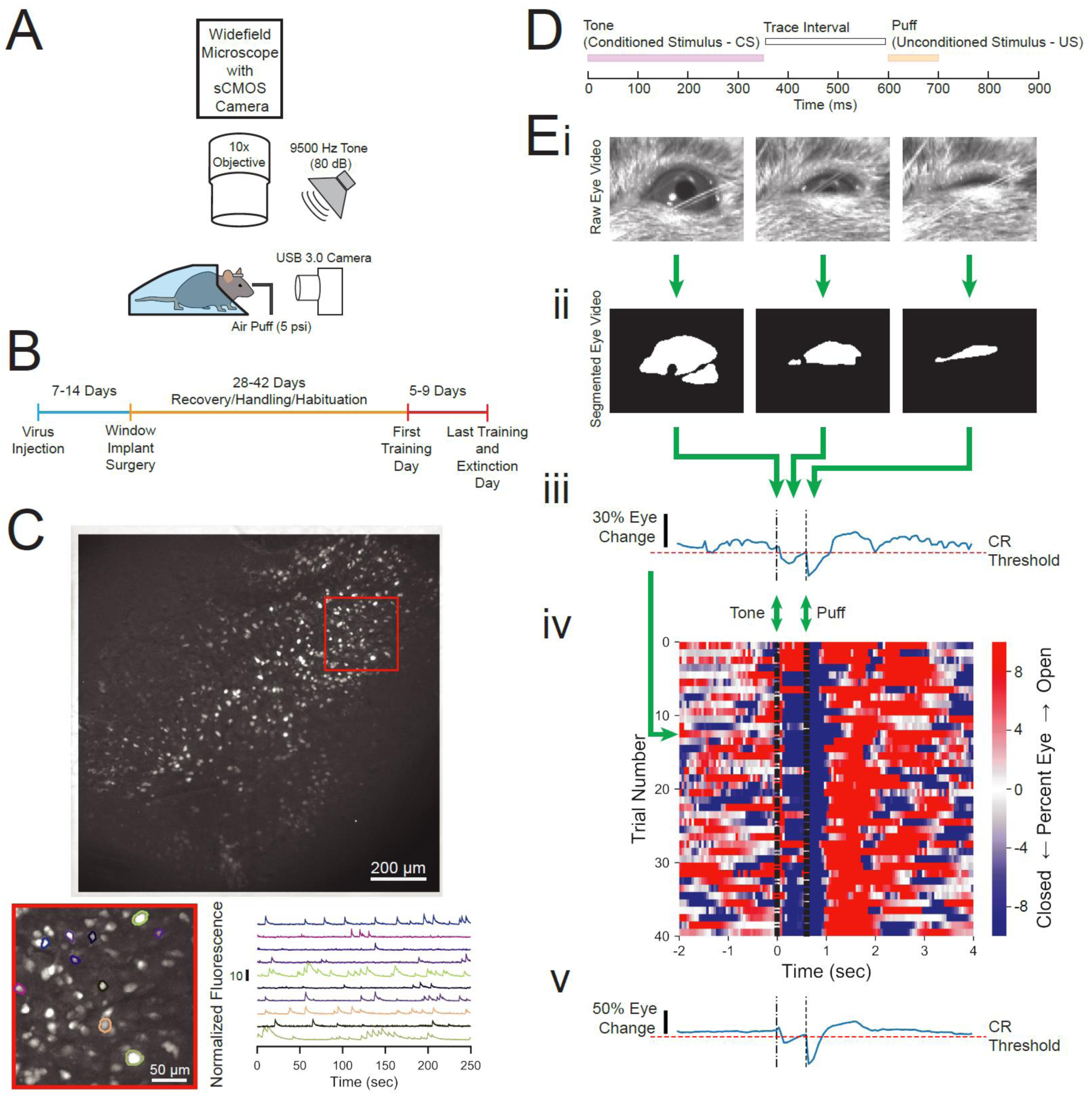
Experimental design and measurement of animal behavior. **(A)** Imaging and behavioral setup. The imaging setup consisted of a microscope with a sCMOS camera, standard wide-field fluorescence optics, and a 10x long working distance objective to image a head-fixed mouse. For the behavioral paradigm, a speaker was positioned near the mouse and a cannula for directing an air puff was placed in front of the right eye. Eye responses were monitored using a USB 3.0 Camera. See methods for full details. **(B)** Experimental timeline. Each animal was injected with AAV-Syn-GCaMP6f and allowed 1-2 weeks for virus expression before surgical window implantation above CA1. The first training day was 4-6 weeks after surgery, and animals were trained and imaged for 5-9 days. **(C)** Full field of view and selected extracted traces. Maximum-minimum projection for one motion corrected video to show example field of view of several hundred cells. Inset: Several selected cells and their corresponding normalized fluorescence trace recordings. **(D)** Within trial design. Trials consisted of a 350 ms tone as the conditioned stimulus (CS), followed by a 250 ms trace interval with no sound, followed by a 100 ms puff of air to the eye as the unconditioned stimulus (US). **(E)** Video eye monitoring and segmentation. **(Ei)** Raw eye frames aligned to the CS, trace interval, and US windows shown above. **(Eii)** Eye frames after segmentation using Fiji^56^ and MorphoLibJ^57^. **(Eiii)** Extracted eye trace for one trial. Eye movement thresholds were calculated for each recording and used to classify whether the mouse eyelid moved enough to constitute a conditioned response. CR – conditioned response. **(Eiv)** Eye trace for all 40 trials of a first training session from one example mouse. Red indicates eye opening, while blue indicates eye closure. Most trials show a conditioned response in blue after the tone but before the puff, and there is a strong blue band after the puff on every trial. **(Ev)** Extracted eye trace averaged over all trials shown in Eiv.

Behavioral response was quantified by segmenting the eye videos and averaging each frame to calculate a temporal trace of eyelid movement (Figure 1Ei-iii). A movement threshold was calculated for each eye trace, fit to a uniform distribution equal to the average eye size. This thresholding method provided a consistent evaluation of behavioral response for each session across mice: eye closure with an amplitude above the threshold between the tone onset and puff onset (tone-puff window) was classified as a conditioned response (Figure 1Ei-Eiv). Using this method, we were able to track the strength of the response to the CS, as well as the strong, persistent eye closure in response to the aversive US on each trial (Figure 1Eiv). This method also allows for consistent calculation of conditioned response within the training session (Figure 1Eiv-1Ev) and across days (Figure 2A). Behavior was measured and scored according to this metric across mice. Animals consistently showed more conditioned responding in the final 20 trials of the last training session (70 ± 11%, mean ± s.d.) compared to the first 10 trials from the first training session (49 ± 26%, mean ± s.d., p=0.0278, one-tailed paired t-test, alpha<0.05 after Benjamini-Hochberg procedure). Additionally, animals consistently exhibited decreased conditioning responding during the final 20 trials of the extinction session (45 ± 21%, mean ± s.d., p=0.0134, one-tailed paired t-test, alpha<0.025 after Benjamini-Hochberg procedure) (Figure 2A-B).

**Figure 2.**
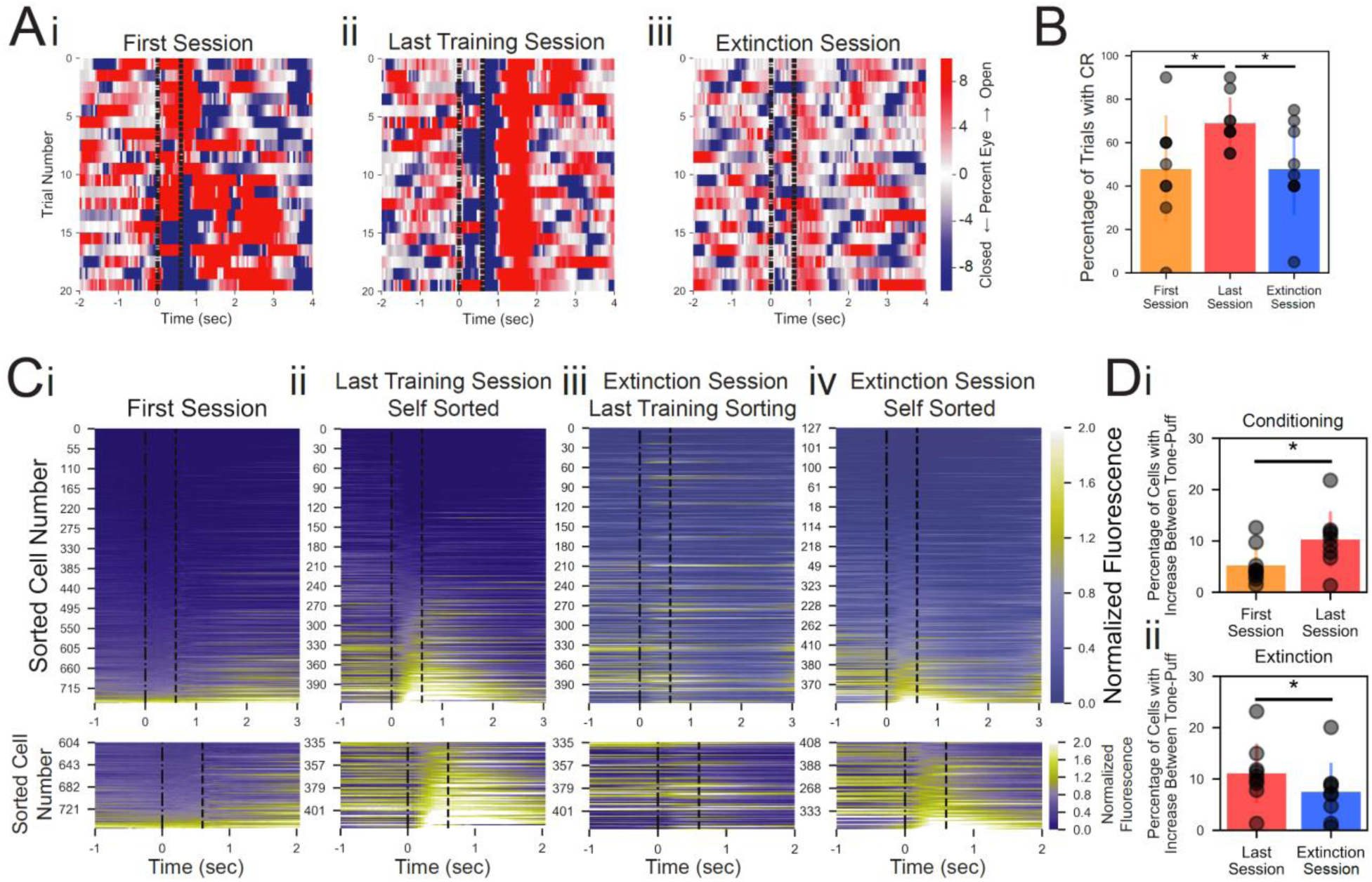
Conditioned responses and neuronal calcium responses increase during conditioning and decrease during extinction. **(A)** Extracted eye traces across days. Red indicates eye opening, while blue indicates eye closure. **(Ai)** Eye trace for 10 tone-only trials and the first 10 trials of the first training session from the same example mouse in Figure 1E. Note the absence of a strong blue band at the time when the puff would normally occur in trials 1-10. Note the absence of blue (indicating absence of conditioned response) during the tone-puff window of the first few training trials (beginning trial 11). **(Aii)** Eye trace for all 20 trials of the last training session for the same example mouse. Note the blue band (indicating conditioned response) during the tone-puff window on most trials throughout the session. **(Aiii)** Eye trace for the last 20 trials of the extinction session for the same example mouse. Note the lack of conditioned response during the tone-puff window on most trials, and note the absence of a strong blue band at the time when the puff would normally occur (as puff is removed during extinction). **(B)** Quantification of conditioned response for all subjects during the first 10 trials of the first session, last 20 trials of the last session, and last 20 trials of the extinction session. Conditioned response rate increased between the first and last training sessions, indicating learning among the mice. Conditioned response rate decreased between the last training session and the extinction session. CR – conditioned response, *p=0.0278 and alpha<0.05 for first vs last session, and p=0.0134 and alpha<0.025 for last vs extinction session, one-tailed paired t-test and Benjamini-Hochberg procedure. **(C)** Trial-averaged calcium recordings. **(Ci)** Top: Trial-averaged recordings sorted by average fluorescence between the tone and the puff for the first training session from an example mouse. Bottom: Zoomed inset of bottom 20% of cells for first training session. **(Cii)** The same as in Ci, for the last training session from the same mouse, sorted by average fluorescence between the tone and the puff for the last training session. More cells showed a consistent trial-averaged response for the last training session than during the first session, as seen by the green bands between tone and puff on the last training session. **(Ciii)** Trial-averaged recordings (plotted as in Ci) of the extinction session, but cell sorting was maintained from the last training session to look at the spatially matched cells. Maintaining sorting revealed a reduction in response to the CS in previously CS-responsive neurons. **(Civ)** The same data as shown in Ciii, but resorted according to the fluorescence between the tone and the puff for the extinction session. Resorting on the extinction session alone shows a new population of cells responsive to CS during the extinction session. **(Di)** Quantification of the proportion of cells responsive to the CS from the first session (yellow) and the last training session (red), *p=0.0463, paired one-tailed t-test. **(Dii)** Quantification of the proportion of cells responsive to the CS from the last training session compared to the extinction session (blue), *p=0.016, paired one-tailed t-test.

### Calcium dynamics in CA1 reflect the behavioral responses during trace conditioning

Imaging CA1 during trace conditioning allowed us to evaluate how activities of large neuron populations (324 ± 246 cells from n=8 mice, mean ± s.d.) are modulated between the first and final day of conditioning training. Furthermore, when extinction was introduced, we imaged the same neurons during trace conditioning and extinction learning, enabling us to investigate whether conditioning and extinction recruit unique cell populations or repurpose the same population. In order to assess the activity of individual cells, imaging sessions underwent several stages of processing. Videos were motion corrected, a projection image was generated across each video, cells were segmented using a semi-automated software, and fluorescence traces for each cell were extracted and normalized for each imaging session (Figure 1C).

A distinct pattern of neuronal responses within the CA1 emerged after multiple days of trace conditioning (Figure 2Ci-Cii). Neuronal responses for each cell were averaged across all trials, and the entire population was sorted by average response intensity during the time period between the tone onset and puff onset (tone-puff window). During the last training session, after multiple days of conditioning, substantially more neurons (11.3 ± 6.2%: mean ± s.d) exhibited an increased calcium response between the tone and puff, compared to the first day of training (5.6 ± 3.7%: mean ± s.d., p=0.0463, one-tailed paired t-test, Figure 2Ci-ii,D), suggesting enhanced CA1 recruitment across training sessions. The percentages of cells that responded on the first and last sessions of training were both significantly greater than the percentages expected by chance (bootstrapped estimation, both: N=1000, p=0.026 & 0.016 for first and last session respectively, one-tailed bootstrap, alpha=0.05, Supp. Figure 1A). These findings suggest that CA1 neurons begin to encode the CS on the first day of training, and the number of neurons encoding the CS increased over several days of conditioning training.

### Extinction learning rapidly recruits new CA1 neurons

The last imaging day included a CS-US training session (referred to as “last training session”) immediately followed by 40 CS-only trials (“extinction session”), allowing all cells to be matched between the two sessions. Behavioral analysis revealed that conditioned responding was significantly reduced as a result of extinction training (Figure 2B). When neuronal responses were averaged together and analyzed as described above, we found that 8.3 ± 5.4% (mean ± s.d.) of cells were responsive to CS during extinction session, which is significantly higher than the percentage expected by chance (bootstrapped estimation, extinction: N=1000, p=0.011, one-tailed bootstrap, alpha=0.05, Supp. Figure 1B).

We compared the responses of individual cells to the CS during conditioning trials and extinction trials by plotting the neuronal responses of the entire population during the extinction session, but maintaining the sorting determined from the last training session. Interestingly, we discovered that most (78.2 ± 15.7%: mean ± s.d.) of the responsive cells during conditioning were no longer responsive during the extinction session (Figure 2Cii-iii). Further analysis of the responses during the extinction session, but resorting the population based on the average GCaMP6f fluorescence intensity during the tone-puff window of the extinction session itself, revealed another population of neurons that were responsive during extinction (Figure 2Civ). However, these cells were a largely novel population of cells that were not responsive to the tone during the previous conditioned learning, as most (67.2 ± 22.3%: mean ± s.d.) of the cells responsive during extinction exhibited no response on the last day of training (Figure 2Ciii-iv). CS-responsive neurons from the extinction session represented a statistically smaller proportion of the population relative to the CS-responsive cells from the last training session (8.3 ± 5.4%: mean ± s.d. versus 11.3 ± 6.2%: mean ± s.d.: p=0.016, one-tailed paired t-test, Figure 2D).

The population of cells that responded to the CS (increased fluorescence during the tone-puff window) during the last training session were termed conditioned (CO) cells. In contrast, extinction (EX) cells were determined as those that responded to the CS during extinction. These populations were largely discrete, as only 21.8 ± 15.7% (mean ± s.d.) of CO cells were also EX cells, and similarly only 32.8 ± 22.3% (mean ± s.d.) of EX cells were also CS cells. These results suggest that during extinction learning, neurons are recruited to encode tone presentations rapidly and emerge in less than 40 trials. Additionally, we have identified two largely distinct populations of neurons that respond to the tone during either conditioning or extinction learning.

### Temporally and spatially distributed populations of neurons encode learning for either trace conditioning or extinction

Because calcium events are sparse, we next considered the reliability of individual cell activation to the tone under both conditioned and extinction learning. Some cells showed responses to the CS on multiple conditioning or extinction trials (example CO cells: Figure 3A; example EX cells: Figure 3B). However, most individual CA1 neurons exhibit highly diverse responses to the tone. CO cells responded to a highly variable number of trials during the last training session (5-78% of trials across all neurons from all mice), with a mean of 11.8 ± 4.9% (mean ± s.d.). Similarly, EX cells responded to 5-63% of trials during the extinction session, with a mean of 10.1 ± 2.7% (mean ± s.d.) of trials during extinction training (Supp. Figure 2). These results indicate that for both conditioning and extinction, most neurons of the entire population contributed to encoding only about 10% of total trials in the session.

**Figure 3.**
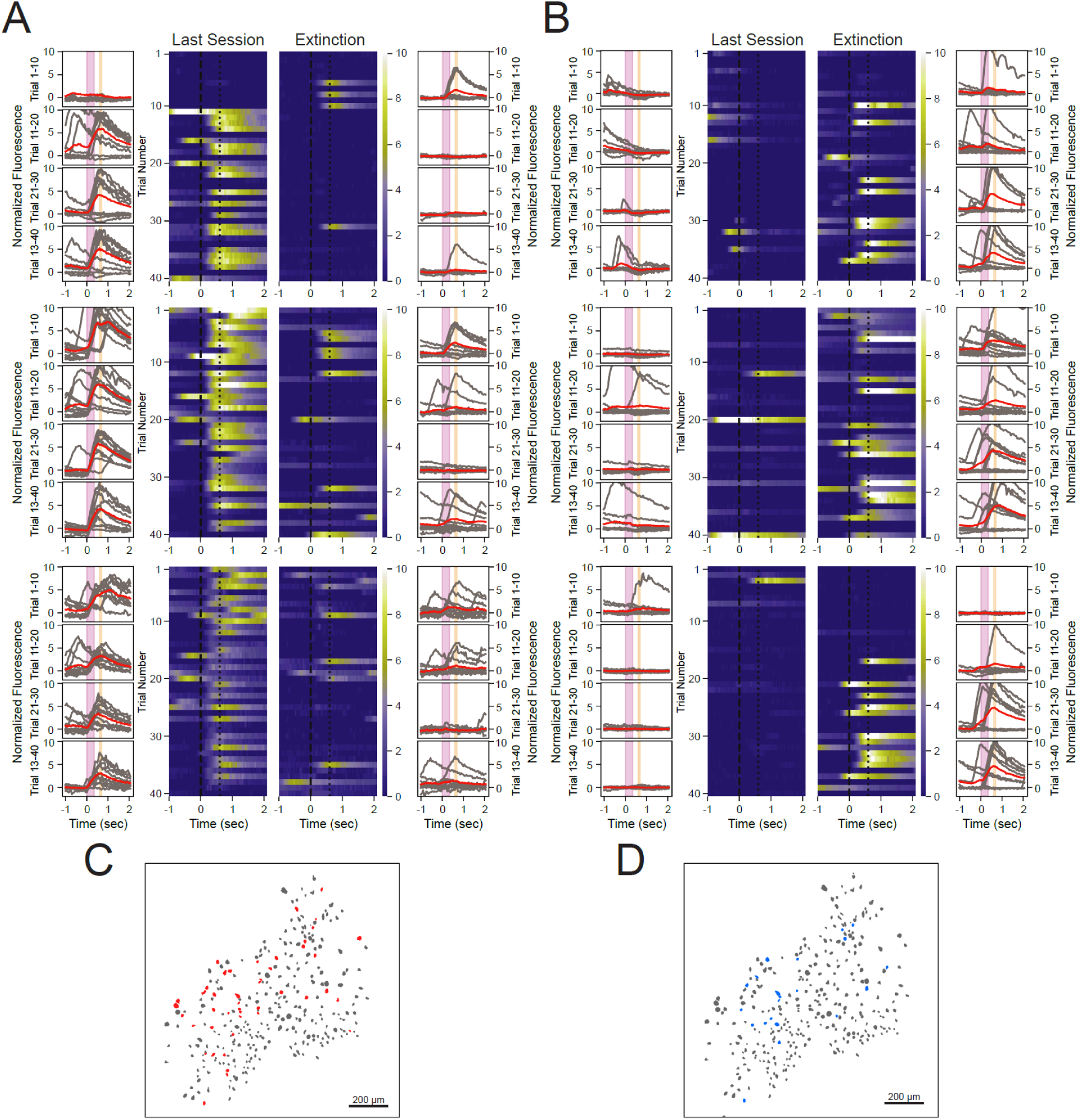
Some neurons show robust responses to the tone during either conditioning or extinction learning, but not both. **(A)** Responses across all trials for three neurons that show a reliable responding during the last training session, termed CO cells. Outer columns are individual trials shown in gray and the average of each 10 trials shown in red. The pink box corresponds to the tone and the orange box corresponds to the puff. Heat maps in the center show each trial for a 3-second time window surrounding the tone and puff presentations. Note some responses of these cells on early extinction trials, but much fewer on later extinction trials. **(B)** The same as in A, for three neurons that exhibit a reliable responding for extinction trials, termed EX cells. Note the lack of response of these cells to the tone during the last training session. Also, note that both the CO and EX cells do not respond on every relevant trial, but show consistent responding across the full session. **(C)** Spatial maps of all neuron masks from a representative animal, with CO cells in red and all other cells in gray. **(D)** The same map as in C, with EX cells in blue and all other cells in gray.

Comparing the spatial distributions of cells suggests that both CO and EX cells are not significantly clustered near one another than would be expected by a random distribution of cell types in CA1 (Figure 3C,D). 3.0 ± 2.5% (mean ± s.d.) of cells within a 100 µm radius of CO cells were other CO cells, which was not significantly different from that expected by random shuffling of cell identity (shuffled=1.9%, N=1000, p=0.17, one-tailed bootstrap, alpha=0.05). 1.6 ± 1.5% (mean ± s.d.) of cells within a 100 µm radius of EX cells were other EX cells, which was also not significantly higher than expected by a random shuffling of cell identity (shuffled=1.3%, N=1000, p=0.24, one-tailed bootstrap, alpha=0.05). These analyses reveal that individual CO cells and EX cells respond on a sparse subset of trials, and that CO and EX cells were heterogeneously distributed within the CA1.

### Co-occurrence analysis reveals differential connectivity between sub-populations of neurons during either trace conditioning or extinction learning

Most CO and EX neurons identified from trial-averaged responses responded on a relatively small percentage of trials, and some cells were so sparsely activated in response to the CS (Figure 4A) that they failed to reach the threshold to be identified as either a CO or EX cell. Given that the majority of individual cells encode CS presentations with low response reliability, network or population responses might more faithfully reveal the role of the CA1 in learning and memory. Thus, we quantified network responses based on pairwise activity of the cell population on a trial-by-trial basis, which summarizes co-activity across all neurons in a “co-occurrence matrix”. While pairwise correlation can give reliable measures over many trials or longer periods of time, the limited number of imaging data points (12) during the short (600ms) tone-puff window of this study made pairwise correlation unsuitable for tracking neuronal calcium responses. The presence of activity during the tone-puff window was determined for each neuron individually by comparing its calcium response between the tone onset and puff onset to an equal time period before the tone. For each trial, if a neuron’s calcium response increased during the tone-puff window, the neuron was assigned a 1, and no increase in activity was assigned a 0 (Figure 4Bi). Taking the outer product of this response vector yielded a co-occurrence matrix of all cell interactions in the population for a single trial based on the response values during the tone-puff window (Figure 4Bii). These single trial co-occurrence matrices were combined on specific subsets of trials and clustered using spectral biclustering to identify neurons that were highly co-modulated for those trials^41^ (Figure 4Bii).

**Figure 4.**
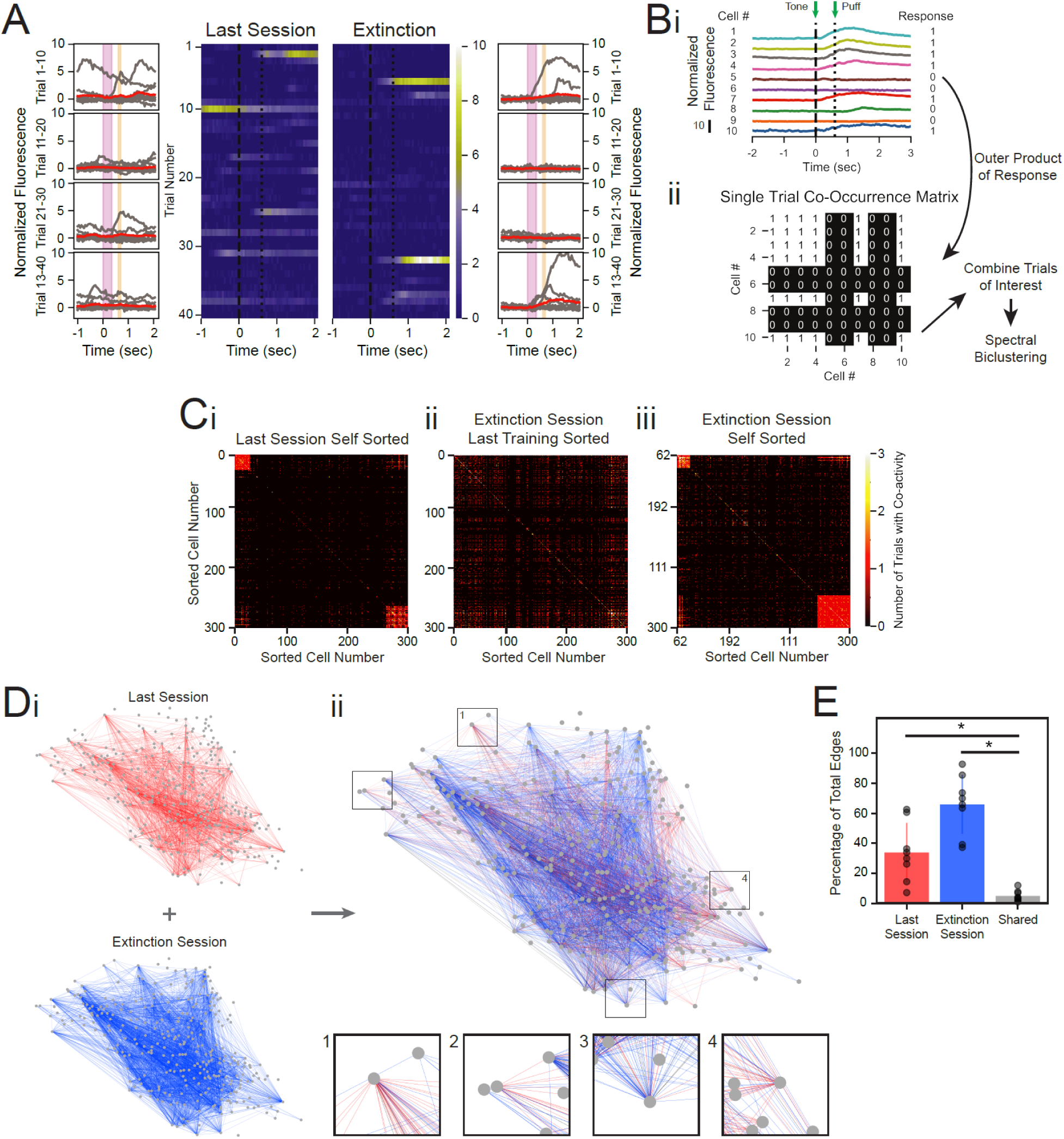
Co-occurrence analysis provides a measure of population activity across individual trials. **(A)** Non-classified cell (neither CO nor EX) that highlights the heterogeneity of responses in the general population of cells, plotted as in Figure 3A and B. **(B)** Schematic of method for constructing single-trial co-occurrence matrices. **(Bi)** A sub-population of cells for one trial that highlights the how the response pattern was determined. If a cell showed an increase in activity during the tone-puff window, compared to the pre-tone period, it was assigned a 1. **(Bii)** The outer product was taken of the vector of responses across the population with itself to generate a single trial co-occurrence matrix. This is a binary matrix where if the ith and jth cells both increase during the tone-puff window there is a 1, but a 0 otherwise. These individual trials can be combined as specific trials of interest, and clustered with spectral biclustering to identify neurons with the highest degree of co-activity across trial types. **(C)** Representative co-occurrence matrices. **(Ci)** Clustering based on co-occurrence matrix for the last 20 trials of the last training session from an example mouse. Note the clusters of cell pairs with high co-activity during the last session. **(Cii)** Co-occurrence matrix for all trials of the extinction session, with sorting maintained from the last training session matrix. Maintaining sorting shows a lack in co-activity between the same cell pairs that were present in the co-active clusters of the last training session. **(Ciii)** The same data as shown in Cii, re-clustered. Re-clustering the extinction matrix reveals separate populations of cell pairs that are highly co-active during the extinction session. **(D)** Connectivity maps created from co-occurrence matrices. **(Di)** Connectivity maps created from the last training session co-occurrence matrix shown in Ci (top) and extinction session co-occurrence matrix shown in Cii and Ciii (bottom). **(Dii)** Last training and extinction session network maps, overlaid. Edges from the last training session are shown in red, edges from the extinction session in blue, and edges present during both sessions are shown in black. Insets: Zoom-ins of four nodes. Note that there are very few edges that overlap (colored black) between the two behavioral conditions. **(E)** Quantification of the proportion of total edges of the last training and extinction sessions that were present during last training session (red), extinction session (blue), or both (gray). p=0.0550 for last vs extinction sessions, *p=0.0023 for last session vs shared and p=0.0001 for extinction session vs shared, paired two-tailed t-test.

Co-occurrence matrices were first generated for the last training session, and clustering was performed on each matrix to identify sub-populations of neurons that were co-active during the last session (Figure 4Ci). Co-occurrence matrices were then generated for the extinction session, but the sorting within each extinction matrix was maintained from the last training session matrix (Figure 4Cii). Similarly to our results observed with trial-averaged responses, the co-active sub-populations identified during the last training session mostly are not co-active during the extinction session. Re-clustering the extinction session matrix revealed new clusters of highly co-active neuron pairs on extinction trials (Figure 4Ciii).

These findings indicate that the pattern of activity within the entire CA1 population as a network is different between the last session and extinction session. To quantify network connectivity, we anatomically mapped co-activity as edges between co-active neurons (nodes) to generate network maps for the last training session and extinction session (Figure 4Di). We found a non-significant difference in the number of edges present in the last training session versus extinction session (34.0 ± 18.4% vs. 66.0 ± 18.4%: mean ± s.d. of the total edges of the last training and extinction sessions combined, p=0.0550, two-tailed paired t-test, Figure 4E). Additionally, the connectivity density and average degree of the two maps across all mice were not significantly different from one another (p=0.218 & 0.6022 for density and degree respectively, two-tailed independent t-test, Supp. Figure 3). When we overlaid the last training and extinction session maps for individual animals, however, we noticed that the edges were largely distinct (Figure 5Cii-iii), with only 5.1 ± 3.4% (mean ± s.d.) of total edges appearing on both the last training session and extinction session maps across animals (significantly smaller than the total number of edges in both behavioral conditions, p=0.0023 & 0.0001 for last training and extinction sessions versus shared edges respectively, two-tailed paired t-test, Figure 5D). These findings indicate that while largely different pairs of neurons are co-active during the last training session versus the extinction session, the involvement of the overall CA1 network (the connectivity density) remains constant during these two behaviors.

**Figure 5.**
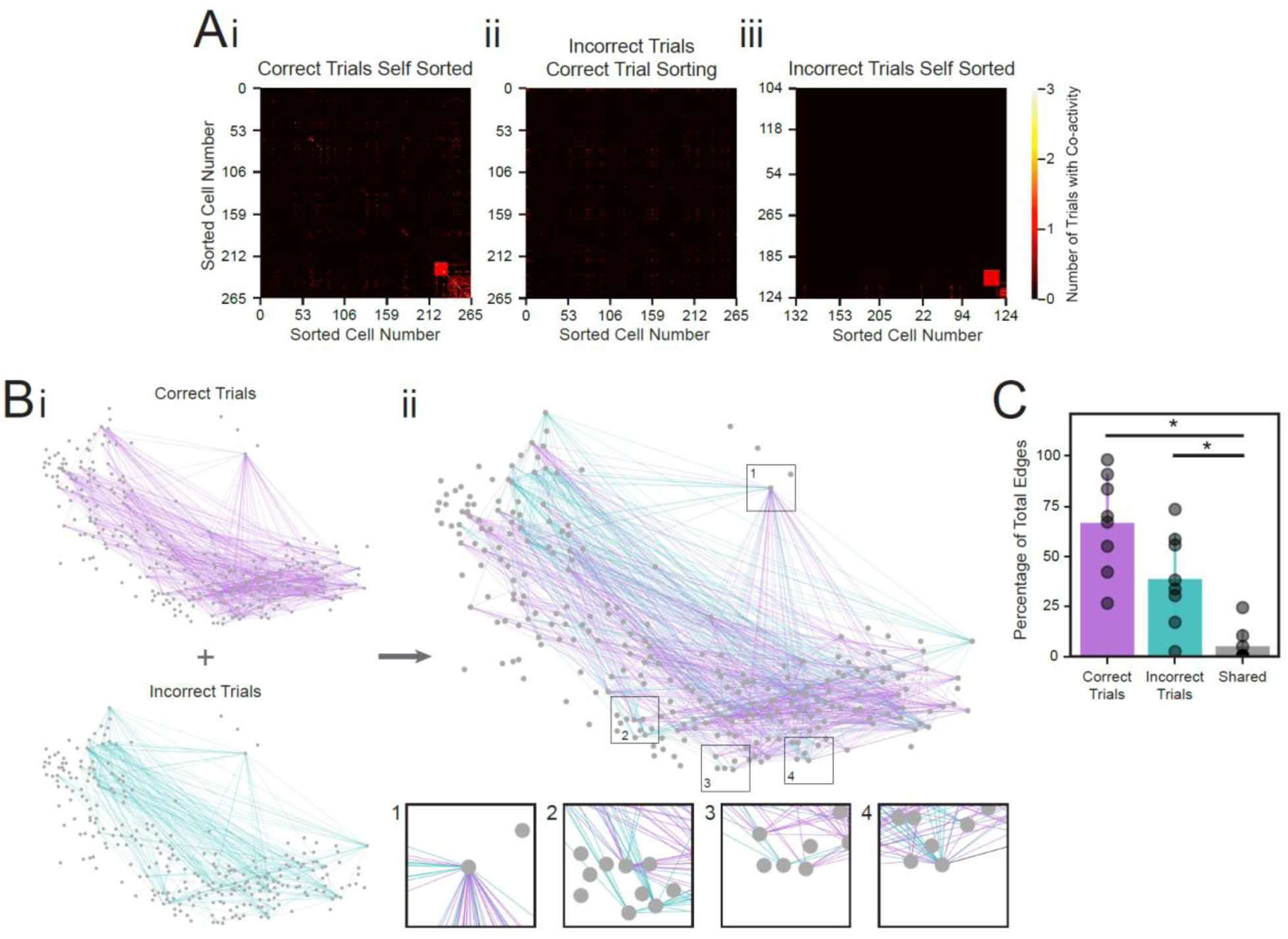
Sub-populations of neuron pairs are differentially activated on trials with different behavioral responses. **(A)** Behaviorally-relevant co-occurrence matrices. **(Ai)** Clustering based on co-occurrence matrix of last training session trials on which the animal performed the correct behavioral response, for a representative animal. Note the clusters of highly co-active cell pairs. **(Aii)** Co-occurrence matrix for the trials from the last training session with the incorrect behavioral response for the same mouse. Sorting is maintained from the last training session to compare co-activity of the same cell pairs between behavioral conditions. Maintaining sorting shows the lack of strong co-activity during incorrect trials in the cell pairs that were highly co-active on correct trials. **(Aiii)** The same data as shown in Aii, re-clustered. Re-clustering reveals new populations of cell pairs that are highly co-active on trials with the incorrect behavioral response. **(B)** Connectivity maps created from co-occurrence matrices. **(Bi)** Connectivity maps created from the correct trials co-occurrence matrix shown in Ai (top) and incorrect trials co-occurrence matrix shown in Aii and Aiii (bottom). **(Bii)** Correct and incorrect trial network maps, overlaid. Edges from correct trials are shown in purple, edges from incorrect trials in teal, and edges present during both trial types are shown in black. Insets: Zoom-ins of four nodes. Note that there are very few edges that overlap (colored black) between the two behavioral conditions. **(C)** Quantification of the proportion of total edges of the last training session that were present during correct trials (purple), incorrect trials (teal), or both (gray). p=0.1355 for correct vs incorrect trials, *p=0.0001 for correct trials vs shared and p=0.006 for incorrect trials vs shared, paired two-tailed t-test.

### Behavioral response during individual trials involves differential connectivity between CA1 neuron pairs

Because the co-occurrence matrix is based on individual trial responses, we used the co-occurrence matrix clustering technique to visualize the co-active neuron populations during “correct” versus “incorrect” trials based on the animal’s behavioral response in the last conditioning session. A correct trial was one in which the animal exhibited a conditioned response, and an incorrect trial was one in which the animal failed to produce a conditioned response. We first generated a co-occurrence matrix for correct trials only, and clustering this matrix revealed sub-populations of neurons that were co-active during correct trials (Figure 5Ai). We then generated a co-occurrence matrix for incorrect trials, but maintained the sorting of the correct trials matrix (Figure 5Aii). Most of the neuron pairs that were co-active during correct trials were not co-active during incorrect trials. Re-clustering the incorrect trials matrix reveals separate sub-populations of cells that were co-active on incorrect trials (Figure 5Aiii).

We generated network maps for both correct and incorrect trial co-occurrence matrices to again investigate differences in activity patterns (Figure 5Bi). The correct trials map included 66.6 ± 22.8% (mean ± s.d.) of the total edges present during the last training session and the incorrect trials map included 38.7 ± 21.6% (mean ± s.d.) of these total edges (p=0.1355, two-tailed paired t-test, Figure 5C). The connectivity density and average degree of the incorrect trials network map were slightly smaller than those for the correct trials map, but not significantly (p=0.2258 & 0.2849 for density and degree respectively, two-tailed independent t-test, Supp. Figure 4). However, the edges were again almost completely distinct (Figure 5Bii-iii). In fact, only 5.3 ± 8.0%: (mean ± s.d.) of total edges appear on both the correct and incorrect trial maps (p=0.0001 & 0.006 for correct and incorrect trials versus shared edges respectively, two-tailed paired t-test, Figure 5C). These findings again highlight the diverse co-activity among neuron pairs in the same behavioral task, but across different behavioral responses, while maintaining the overall CA1 network activation.

## Discussion

In this study, we provide the first detailed, real-time evidence that largely distinct populations of neurons within the hippocampus respond to a trace conditioned stimulus during either conditioned learning or extinction learning. It has been previously reported that two functionally distinct neuron populations are activated by fear conditioning and extinction in the amygdala^31^. A subsequent study looking at the CA1 region of the hippocampus in a contextual fear paradigm revealed distinct changes in gene phosphorylation states in fear conditioning and fear extinction in largely non-overlapping neural populations^32^. However, in this study, cFos and pERK immuno-activity were used as markers of conditioning and extinction, and were measured hours to days after the respective training. These results provided the first important insight into the potential for distinct population encoding, but the indirect nature of the activity markers and the time course for immuno-quantification does not allow for the distinction between rapid or gradual evolution of conditioning and extinction neuron populations. Further, these markers could be dependent on new protein synthesis or long-term plasticity, and are not measures of the dynamic learning process that occurs during conditioning and extinction.

In order to better understand the dynamic relationship between conditioning and extinction learning in the hippocampus, and to further investigate whether distinct populations encode these learning events, we used calcium imaging to monitor the activity of individual cells throughout conditioning and extinction learning paradigms. We applied trace conditioning because (1) it lends the advantage of a singular defined stimulus to which neural activity can be easily aligned and measured between the two different paradigms, (2) it allows training for both conditioning and extinction to occur in the same imaging session, and (3) learning during trace conditioning evolves over multiple trials, unlike fear-based paradigms where learning often occurs over very few trials. Interestingly, we found that rate of conditioning was highly animal-dependent, evolving gradually in most animals and rapidly in a subset of mice that showed substantial responding to the CS within 15-20 trials. Although acquisition of trace eye-blink conditioning can occur over dozens of trials in a single session^42^, most of our animals showed gradual acquisition, strongly reflected in the increase in proportion of neurons that were active during the tone-puff window from the first day to the final, and in the increase in conditioned response rate after multiple days of training. Overall, our results support the idea that robust conditioned learning gradually evolves over many days as new neurons are recruited to encode the stimulus, and reflect previous electrophysiology studies in rabbits and rats where the time course of learning is slow and evolves through many CS-US pairings^43,44^. In contrast, extinction learning was rapid across all subjects; a new population of neurons that responded to the now extinguished tone emerged within one session of forty trials. Previous work has implicated the prefrontal cortex and septal cholinergic inputs to be critical to the process of extinction, and these pathways may play a pivotal role in the rapidity of extinction neuron emergence^45–47^. More work will need to be done on this front to determine whether reducing or silencing these inputs could delay or block the emergence of extinction-selective neuron populations. In addition, it is possible that extinction learning can occur rapidly because a meaningful memory schema already exists. Studies probing updates in location encoding of familiar places suggest memory re-encoding can occur rapidly for spatial information^48,49^, and our observation for extinction learning may reflect a manifestation of this principle. Finally, EX neurons were found embedded in the pyramidal cell layer and were undistinguishable from CO neurons in terms of their activity profile. This finding suggests that EX neurons are likely excitatory pyramidal cells, which is consistent with the description of extinction-relevant neurons in the amygdala and p-ERK+ extinction cells in the hippocampus as excitatory neurons.

Given the likelihood that EX cells are excitatory pyramidal cells, it is unlikely that EX neurons inhibit CO neurons directly, but may instead mediate the activity of interneurons that have a prominent role in suppressing CO neurons as extinction training evolves. Since EX activity can emerge rapidly, the mechanism of interaction between CO and EX activities may be an important future direction that could benefit the treatment of anxiety-based disorders such as post-traumatic stress disorder (PTSD), which is characterized by over-generalization of fearful stimuli to neutral contexts and impairments in development of extinction learning^50–55^.

Calcium imaging is a powerful tool to understand how large populations of neurons function at the population level. However, when investigating dynamic or rapid network changes, as in extinction learning, it can be difficult to decode the information present in the population using traditional analysis techniques. For example, we had low confidence in the pairwise correlation values measured over the brief tone-puff window (600ms, 12 data points) on a trial-by-trial basis. Traditional single-trial analytic approaches usually cannot find meaningful correlations with such limited data. Thus, our development of a co-occurrence-based approach provides a robust way to measure and observe the trial-by-trial evolution of the population responses, and a means to assess some cells’ contributions that might be otherwise overlooked, or overstated, in trial-averaged data. Additionally, it can be used to break down and compare trials by specific behaviors (i.e., correct versus incorrect) or other variables that may change across trials over time. Finally, the co-occurrence matrix allows us to consider connectivity maps of entire neuron populations, an intuitive way to visualize and investigate the patterns of neural activation. Overall, the co-occurrence matrix is a useful technique for monitoring the evolution of population responses over time from high-dimensional calcium imaging datasets.

Using the co-occurrence matrices, we found that CA1 neurons’ connectivity changes drastically between conditioned learning and extinction training, but also between trials with the correct or incorrect behavioral response during conditioned learning. While some cell pairs may participate in both learning conditions, pairs of neurons are differentially activated during different types of learning, indicating a role of network activation and response in the CA1.

Because the co-occurrence matrix provides information at single trial level, it allows the examination of variations between individual trials. In our analysis of correct and incorrect trials, we found that on any given trial, a very small number of neurons were active, indicating that trial encoding may depend on the contributions of large populations of sparsely active neurons. This provides enlightening information about how the hippocampus may represent and encode behaviorally relevant stimuli. Our results indicate that on each trial only a subset of the appropriate sub-population is activated to encode the relevant features of that trial, but also that these subsets work together to create a larger network that represents learning across an entire session, which may be critical to the encoding or retrieval of learning and memory in CA1.

## Materials and Methods

### Animal Surgery and Recovery

All animal procedures were approved by the Boston University Institutional Animal Care and Use Committee. A total of 9 female C57BL/6 mice, 8–12 week old at the start of the experiments, were used in these studies (Taconic; Hudson, NY). To estimate sample size, power analysis was based on effect size differences found in our previous trace conditioning calcium results recorded in the hippocampus^33^. Power analysis was performed using G*Power 3.1.9.6 (http://www.gpower.hhu.de), applying a one-tailed Wilcoxon signed-rank test utilizing α = 0.05 and a power (1-β) value of 0.80. Following arrival from the vendor, mice were allowed to habituate to the vivarium for 2+ weeks prior to surgery. Animals were group housed during this time. Animals first underwent viral injection surgery targeting the hippocampus under stereotaxic conditions (AP: −2.0 mm, ML: +1.5 mm, DV: −1.6 mm). Mice were injected with 250 nL of AAV9-Syn-GCaMP6f.WPRE.SV40 virus obtained from the University of Pennsylvania Vector Core (titer ∼6e12 GC/ml). All injections were made via pulled glass pipettes (diameter: 1.2 mm) pulled to a sharp point and then broken at the tip to a final inner diameter of ∼20 μm. Virus was delivered via slow pressure ejection (10-15 psi, 15-20 ms pulses delivered at 0.5 Hz). The pipette was lowered over 3 min and allowed to remain in place for 3 min before infusion began. The rate of the infusion was 100 nL/min. At the conclusion of the infusion, the pipette remained in place for 10 min before slowly being withdrawn over 2-3 minutes. Upon complete recovery (7+ days after virus injection, mice underwent a second procedure for the implantation of a sterilized custom imaging cannula (OD: 0.317 cm, ID: 0.236 cm, height, 2 mm diameter), fitted with a circular coverslip (size 0; OD: 3mm) adhered using a UV-curable optical adhesive (Norland Products). To access dorsal CA1, the cortical tissue overlying the hippocampus was carefully aspirated away to expose the corpus callosum. The white matter was then thinned until the underlying tissue could be visualized through the surgical microscope. The window was then placed and centered above the hippocampus. During the same surgery, a custom aluminum head-plate was attached to the skull, anterior to the imaging cannula.

### Animal Training and Trace Conditioning Behavioral Paradigm

Mice were trained on a trace eye-blink conditioning task similar to what was described previously^33^. Animals were allowed at least 2 weeks to recover from window surgeries, followed with an additional 2-4 weeks of handling and habituation to being head-fixed underneath the microscope (Figure 1Bii). Each animal received at least 3 habituation sessions prior to the first recording day. Habituation was performed in the dark with the imaging LED illuminated to the same intensity as it would be for recording sessions.

Following habituation, training for the eye-blink conditioning task began. Each trial consisted of a 350 ms long 9500 Hz tone (conditioned stimulus - CS) at 78-84 dB followed by a 250 ms trace interval, followed by a 100 ms puff to the eye (unconditioned stimulus – US) at 4.2-6 psi (Figure 1Bi). The ambient noise level ranged between 55-60 dB. Inter-trial intervals for each presentation were pseudo-randomized within a recording session with an inter-trial interval of 35 ± 5 seconds. The first 20 recording trials consisted of tone only presentation without the puff. Animals were then presented with either 60 tone-puff trials per day for 8 days, or 80 tone-puff trials per day for 4 days. The final recording session consisted of 20 or 40 tone-puff trials as the last learning session, followed by 40 extinction trials, where the puff was removed but the tone continued for those trials. Behavioral stimuli were generated using a custom MATLAB script that delivered TTL pulses for the tone and puff via an I/O interface (USB-6259; National Instruments, Austin, TX). Behavioral TTL pulses and imaging frame timing were digitized and recorded (Digidata 1440A; Axon CNS Molecular Devices, San Jose, CA or RZ5D Bioamp Processor; Tucker Davis Technologies, Alachua, FL) to align behavioral data and imaging frames.

Mouse eye positioning was captured using a Flea3 USB 3.0 camera (FL3-U3-13S2C-CS; Richmond, BC, Canada) and the Point Grey FlyCapture 2 software, after illuminating the eye and surrounding area with an IR lamp positioned approximately 0.05-0.5 meters away from the mouse.

### Wide-field imaging

A custom-built wide-field microscope was used to record neuronal calcium responses during animal learning and behavior as previously described^33^. Briefly, the animal was head-fixed below the microscope on an articulating base (SL20 Articulating Base Ball Stage; Thorlabs Inc, Morganville, NJ) and a custom-machined attachment for the headbar, with the animal being covered by an elastic self-adherent wrap to reduce movement during recording. The microscope consisted of a scientific CMOS (sCMOS) camera (ORCA-Flash4.0 LT Digital CMOS camera C11440-42U; Hamamatsu, Boston, MA) was used in conjunction with standard optics for imaging GCaMP6 and a 10x magnification objective (Leica N Plan 10 × 0.25 PH1 or Mitutoyo Plan Apo Long WD Objective 10 × 0.28). Images yielded a field of view 1.343 mm by 1.343 mm (1024×1024 pixels) and were acquired at a 20 Hz sampling rate and stored offline for analysis.

### Data Analysis

#### Behavior Eye-Blink Segmentation and Analysis

First, each raw video was segmented using Fiji^56^ and the MorphoLibJ plugin^57^ to generate a binary mask video corresponding to the animal’s eye. To do so, each frame of this binary video was summed and normalized by the average eye size to generate a trace corresponding to the percentage of eye closure over time. First, image stacks were loaded as grayscale images, Gaussian filtered with a radius of 2, and thresholded to include only the eye range. Videos were converted to binary, holes were filled, and the Particle Analyzer feature was used to exclude all ROIs on the edges of the videos above the thresholded value. The MorphoLibJ plugin^57^, was used to label connected components with a connectivity of 26. A custom Jython script (StepIntegers.py) was used to determine the connected components that existed across all image frames, which were merged into one connected component. Lastly, to capture any additional smaller connected components that commonly were created at or around the time of blinks, another custom Jython script (FindModalValues.py) was used to capture these remaining components which were then merged into the final connected component. All other connected components not a part of this singular merged component were dropped from the binary mask stack which was saved for eye-blink trace generation.

Eye-blink traces over time were generated by summing the binary pixels corresponding to the segmented eye in each video frame and dividing by the average eye size across the whole video. An eye conditioned response was classified by calculating a threshold of deviation from the standard eye sizing. The threshold was calculated by fitting a line to the central 95 percentile of the full eye-trace, and a deviation of eye size below 2% from this line was classified as a conditioned response. This is equivalent to when the residuals deviated by 2% from a uniform distribution fit to the eye trace that was equal to the average eye size. Each time the eye-trace showed a decrease below this threshold between the tone onset and puff onset, that trial was classified as a conditioned response trial. For comparison of behavioral performance, the final 20 trials of behavioral responding were selected for analysis from the extinction training session and the final training session. Conditioned responding measured from the first session was limited to the initial 10 trials. Trials were divided as such to capture periods of stability within the process of learning, as speed of learning acquisition varied between individual mice.

#### Movement Correction

Motion correction of videos was done using *ptmc*, an open-source, parallel python version (github.com/HanLabBU/ptmc) of an image stabilization process published previously^33^. Briefly, each frame was motion corrected by median filtering each image to remove noise, homomorphic filtering the image for edge detection, and comparing the frame with a reference image to determine how many pixels to shift that specific frame. This process was run in parallel by first motion correcting the first multi-page tiff stack (2047 frames) with to an average projection image of the noisy, non-corrected video. This corrected video stack was used to generate a new reference image that was sent out in parallel with every frame across the whole video, including the first tiff stack used to generate the reference.

#### Neuronal Trace Extraction

After motion correction, regions-of-interest (ROIs) corresponding to cells were selected using a semi-automated custom written MATLAB software called *SemiSeg* (github.com/HanLabBU/SemiSeg). First, a projection image (Max-minus-Min) across the whole video stack was calculated for selecting ROIs. This static frame was loaded into SemiSeg and the full boundary of the ROIs was selected by a user selecting a sub-region of the image that was thresholded to determine the corresponding pixels within that region that correspond to a cell. After all cells were selected from the projection image, pixels corresponding to these ROIs were averaged together spatially to calculate a temporal trace for each neuron.

For sessions where ROIs were matched to one another, spatial ROI maps were co-registered using frame-wise cross-correlation. ROIs were then matched using a greedy method that required the centroid of cells to be within 50 pixels of one another and had to have at least 50% of their pixels overlap, as was published previously^58^. Cells that did not meet both of those criteria were removed from the matched dataset for comparison.

#### Fluorescence Trace Normalization

Each neuron’s fluorescence trace was normalized after a local background subtraction calculated for each trace. A local background trace was calculated by finding the centroid for each ROI, and measuring a circle approximately 10 cell widths in radius (100 pixels) and subtracting the area for the ROI from that circle. The pixels in this local background were averaged together spatially to measure a temporal background trace. Background traces were subtracted from each cell’s measured trace to remove local fluctuations from scattering in wide-field imaging. The baseline calcium level was calculated for each cell by fitting a normal distribution to the lowest 50 percentile of the data and using the mean of this distribution as the baseline calcium level. This baseline was subtracted from each locally corrected trace, and data was scaled by 5% of the maximum range of the full calcium trace.

#### Determination of Increased Activity Cells

For trial-averaged analysis, all trials of the last training session were included. Fluorescence for the 12 data points (600 ms) within the interval between tone onset and puff onset (tone-puff window) was compared to the 12 data points prior to the tone. As cells might have only randomly responded once across all trials during this time window, a threshold was selected to capture cells with regularly occurring or very strong responses to the tone. Thus, a cell was classified as having increased if the average fluorescence during the tone-puff window was larger than the average fluorescence for the pre-tone window by 0.15. This 0.15 value is equivalent to a neuron having a several 5% increases (normalized value of 1) on at least 6 trials on one end of the spectrum, or one large 30% increase (normalized value of 6) on the other end of the spectrum. This was the threshold used for all statistics and for comparison with the network measure. All cells that increased in fluorescence by this 0.15 value during the last training session or extinction session were deemed conditioned (CO) or extinction (EX) cells, respectively.

#### Bootstrapping Procedure

Trial-averaged bootstraps were calculated for each mouse to determine what percentages of cells would be expected to increase in random recordings not tied to the tone-puff learning paradigm. Timing between each pseudo-randomized tone-puff was maintained to account for any periodicity effects during the recording, and bootstraps were calculated by circularly permuting the tone-puff timing across all traces. The number of shuffled permutations performed was 1000 for each mouse. The timing of each new randomized tone-puff was averaged together across all shuffled trials, and the percentage of cells that increased between the tone and puff was measured. The percentage of cells increased for each mouse was averaged together to get a population estimation of the mean number of cells expected to increase within the population, which could be compared to the measured percentage of the population that increased between the actual tone-puff across all trials.

#### Spatial Cell Identity Bootstrapping

Bootstraps of shuffled cell identity distributions were calculated for comparison against the observed distribution of cell identities. A 100 micron radius (76 pixels at 1.312 microns/pixel) was drawn around each cell. The number of segmented cells that existed within that spatial distribution were calculated and a percentage of either CO or EX cells was determined from the cell identities from the cells within that radius. For bootstrapping, the same number of CO or EX cells that was segmented for each recording session were randomly selected and the same calculation within a 100 micron radius was calculated. The measured percentages were then compared to the bootstrapped values for statistical confidence.

#### Co-Occurrence Network Creation

Individual trial co-occurrence matrices were created for each pair of cells across every trial. For consistency in the co-occurrence analysis, the last 20 trials of the last training session were included. For each tone-puff window, the mean value of the 600 ms (12 data points) between the tone onset and puff onset was compared to the 600 ms before the tone. If this value was greater than 1 on a normalized trace, corresponding to 5% the maximum peak value of a trace, then the neuron was labelled as responding. The result was a binary vector of 0s and 1s of length N, where N is the number of cells recorded in the population. The outer product of this vector was taken with itself for the whole population to yield an NxN co-occurrence matrix. This matrix is 1 if both the ith and jth cells were activated between the tone-puff, and 0 otherwise.

Once a co-occurrence network was generated for each trial, they could be combined for further analyses by summing on certain trials of interest. For this analysis, co-occurrence matrices were summed across either the last training session, the extinction session, “correct” trials of the last training session, or “incorrect” trials of the last training session. Once a trial combination map was created, spectral biclustering was performed for a 3×3 cluster pattern using the Python machine learning package scikit-learn^41,59^.

#### Network Map Generation

Anatomical spatial information was combined with the co-occurrence matrix to generate network maps using the Python library NetworkX. The centroid of each ROI was used as the position of the corresponding node, which represent the cells of the imaging session. Since the co-occurrence matrix is symmetric, the lower triangular matrix is used to generate the edges of the network. Co-activity between two cells is represented as an edge between the corresponding nodes. For example, the ith cell and jth cell would be connected by an edge if Ai,j is non-zero, where A is the NxN co-occurrence matrix. The combined map of correct and incorrect edges was created by summing the co-occurrence matrices of correct and incorrect trials of the last training session.

#### Quantification of Network Properties

Percentage of edges in each correct or incorrect network is calculated as:

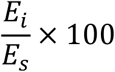

where *E*_*i*_ is total edges in the individual network and *E*_*s*_ is total edges in the session network (comprised of both correct and incorrect trial edges). Percentage of shared edges is calculated as:

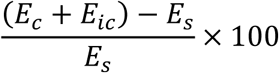

where *E*_*c*_ is total edges in correct network and *E*_*ic*_ is total edges in incorrect network.

Python package NetworkX was used to calculate network density, which is defined as:

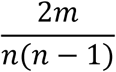

and degree is defined as:

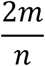

where m is number of edges and n is number of nodes.

## Acknowledgements

K.R.H. and S.S. performed data analysis. R.A.M., H.J.G., and A.I.M. conducted the animal experiments. M.A. and R.K. helped with animal husbandry, habituation, and training. B.N. helped with data analysis. R.A.M., K.R.H., H.J.G., and X.H. wrote the manuscript. X.H. supervised the study.

---

**Supplemental Figure 1.**
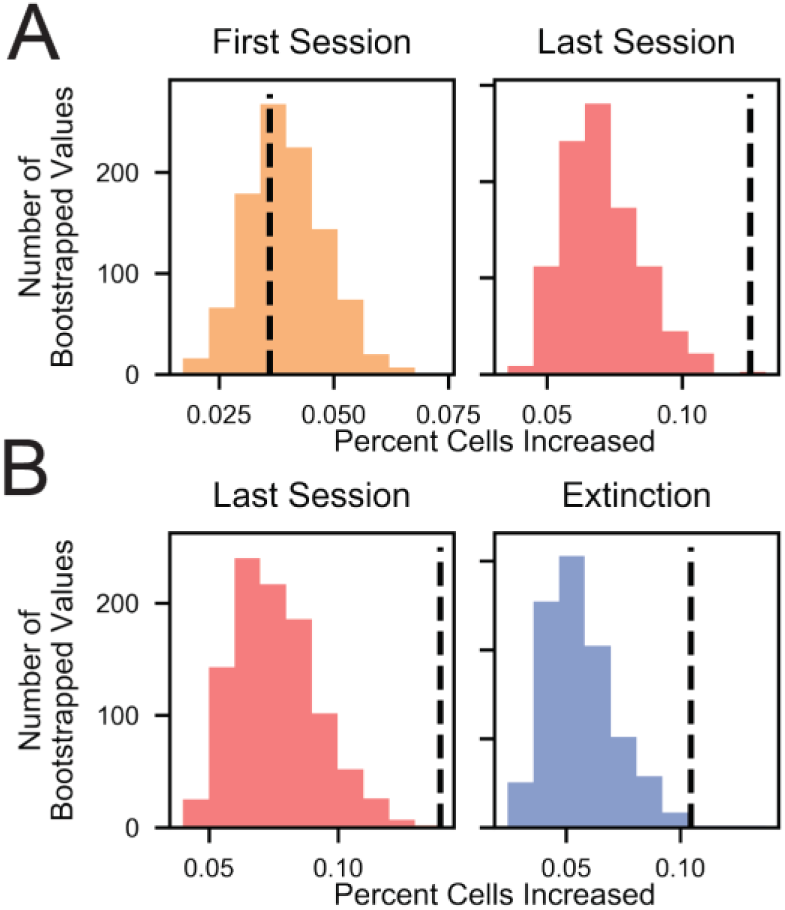
**(A)** For the first session and last training session, bootstrapped distributions of the percentage of cells with an increase between the tone and puff after circularly shuffling the tone-puff locations 1000 times. Dashed black line shows percentage measured with the non-shuffled (experimental) tone-puff locations. **(B)** Bootstrapped distributions comparing the matched last session and extinction session recordings, computed similarly as in A.

**Supplemental Figure 2.**
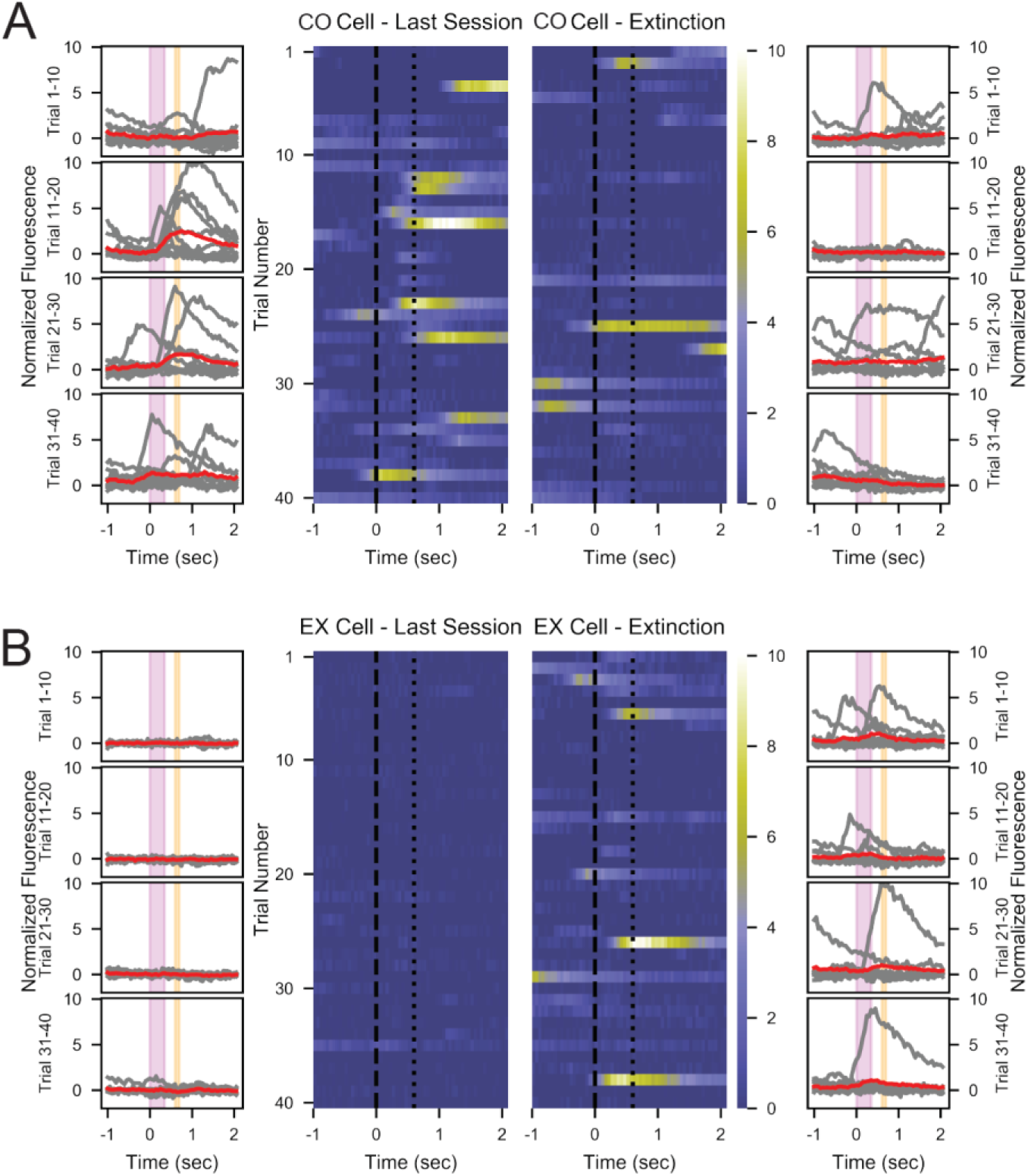
**(A)** Individual single trial responses for CO cell that shows an average level of responding on the last training session. Outer columns have individual trials shown in gray, with the 10-trial average shown in red. The pink box corresponds to the tone interval, and the orange box corresponds to the puff. Center heat-maps show each trial for the window surrounding the tone-puff time. **(B)** The same as in A, for an EX cell that exhibits an average level of responding for extinction trials. Note the lack of response of these cells to the tone during the last training session.

**Supplemental Figure 3.**
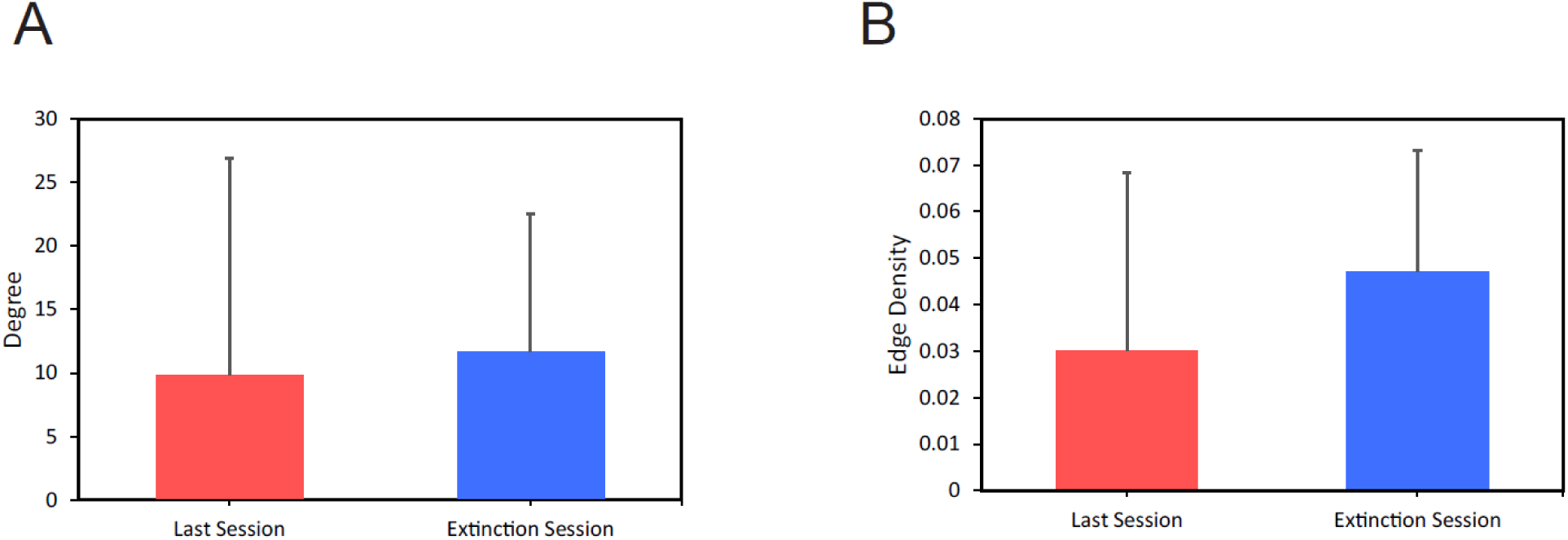
**(A)** Average degree and **(B)** density of connectivity maps for the last training and extinction sessions across all animals, shown mean + s.d. p=0.6022 for average degree and p=0.2018 for density, paired two-tailed t-test.

**Supplemental Figure 4.**
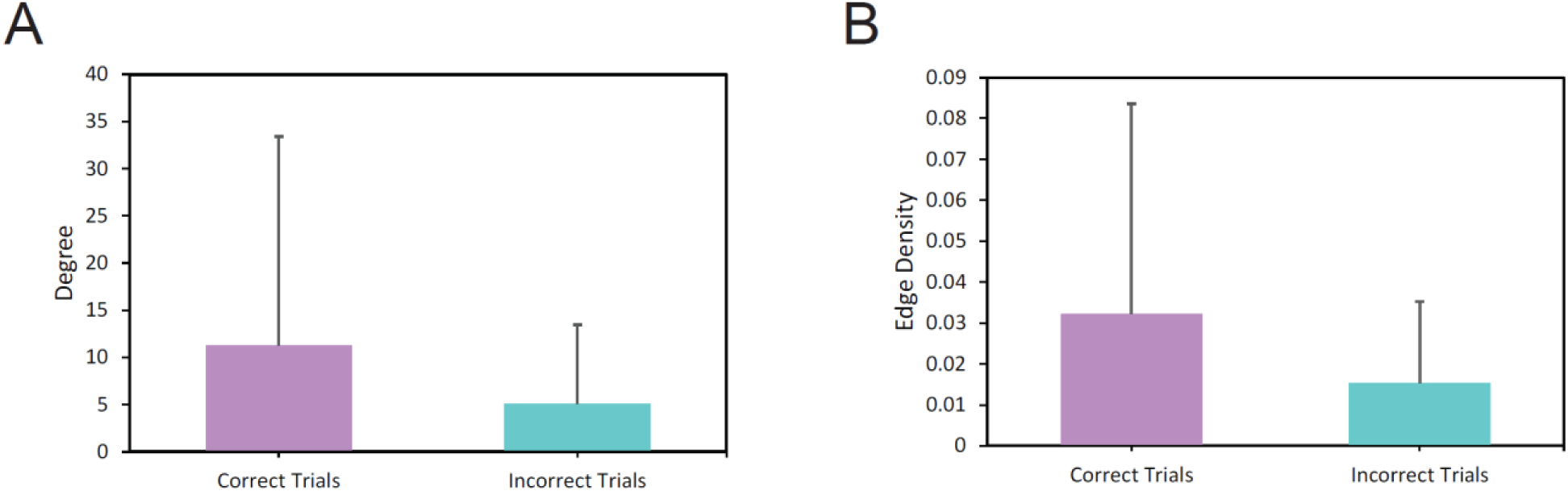
**(A)** Average degree and **(B)** density of connectivity maps for correct and incorrect trials across all animals, shown mean + s.d. p=0.2849 for average degree and p=0.2258 for density, paired two-tailed t-test.

